## Supplemental Figures for "Non-invasive recording from the human olfactory bulb"

---

##### Calculation of time-window of interest

To gain an understanding of what time-window of interest to search within for an olfactory bulb signal, we searched the literature for measures of the olfactory signal latency. Because few measures exist in humans, all values below are based on measurements in model animals and mostly rodent models. Moreover, values below are estimates of the minimal latency time and the upper window is based on actual recordings of neural signal in response to an odor presentation from the human piriform cortex[1-2], an area upstream from the olfactory bulb. These values are estimates and, naturally, the true latency will very between trials, odors, and individuals.

*Estimates of latency within each major processing stage*

Olfactory odor delivery time: 200 ms[3] (*adjusted for in calculations and all figures*)

Mucosa diffusion latency: ~30 ms[4]

Olfactory sensory neuron to first spike: ~30 ms[5]

Conduction delay (~0.2m/s x ~7 mm) ~28 ms[6]

Latency M/T response ~10 ms[7]

Period of interest: **98ms** (set by above estimates, not including odor delivery time) to **300ms** (set by recordings from the human piriform cortex[1-2]).

### Supplementary Material

#### SUPPLEMENTARY FIGURES

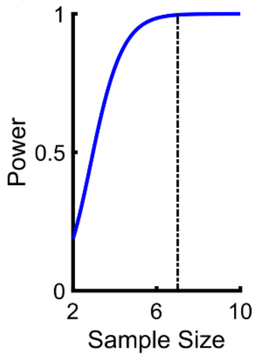


**Supplementary Fig 1.** Estimate of power of the statistical test for the particular hypothesis, and the specific design used, reaches the power of 1 for a minimum sample size of 7 individuals. Power close to 1 means that the probability of rejecting the null hypothesis when H1 is true is large.

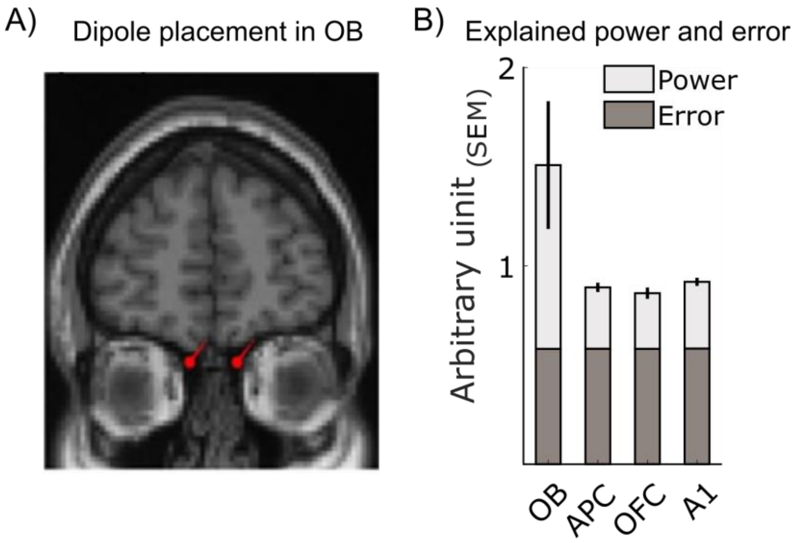


**Supplementary Fig 2. A)** Example of fitting of dipole in bilateral olfactory bulb for guided signal source analyses. **B**) Total explained power of four fitted dipole solutions demonstrating that the OB has the largest explained power. OB=olfactory bulb, APC= anterior piriform cortex, OFC = Orbitofrontal cortex, A1 = primary auditory cortex.

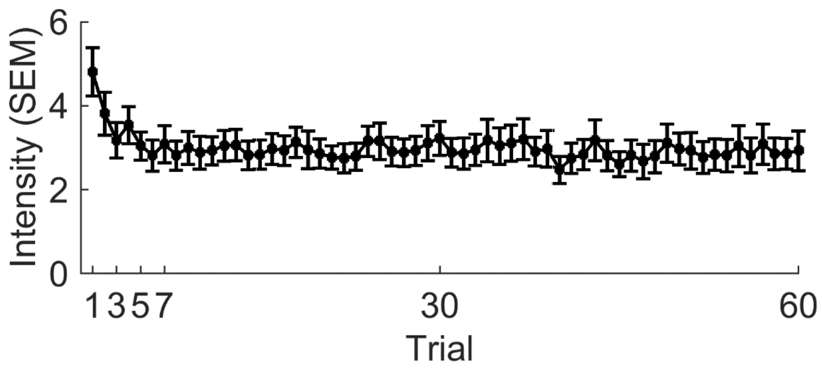


**Supplementary Fig 3.** Average rated perceived intensity across trials. Error bars indicate standard error of the mean (SEM).
